## Supplementary material for "The midbody interactome reveals new unexpected roles for PP1 phosphatases in cytokinesis": This file includes: Supplementary Figs. S1 to S5 Supplementary Tables S1 and S2 Captions for Supplementary Videos S1-S8 and Supplementary Data S1-S5

Supplementary Figs. S1 to S5
Supplementary Tables S1 and S2
Captions for Supplementary Videos S1 to S8
Captions for Supplementary Data S1 to S5

**Other Supplementary Information for this manuscript include the following:**

Movies S1 to S8
Data S1 to S5

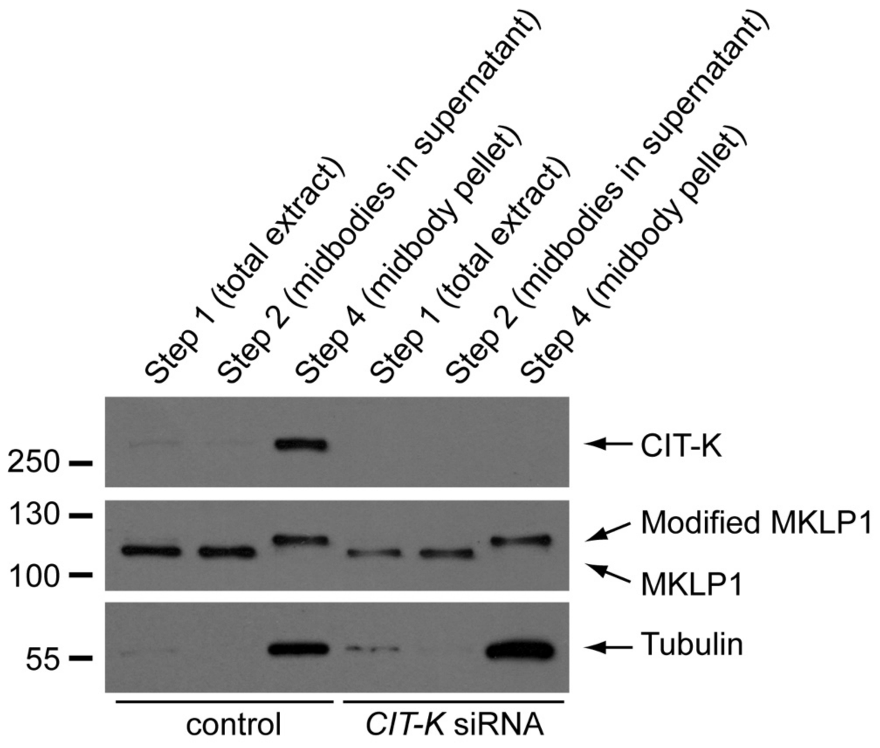


**Supplementary Fig. S1. Purified midbodies are enriched for known midbody proteins**. HeLa cells treated with siRNAs directed against either a random sequence (control) or *CIT-K* were synchronized in telophase to purify midbodies and aliquots from the different purification steps (see Materials and Methods) were analyzed by Western blot to detect CIT-K, MKLP1 and tubulin. The numbers on the left indicate the sizes in kDa of the molecular mass marker. Note that the slower migrating band detected with the anti-MKLP1 antibody is very likely a post-translational modified form of MKLP1.

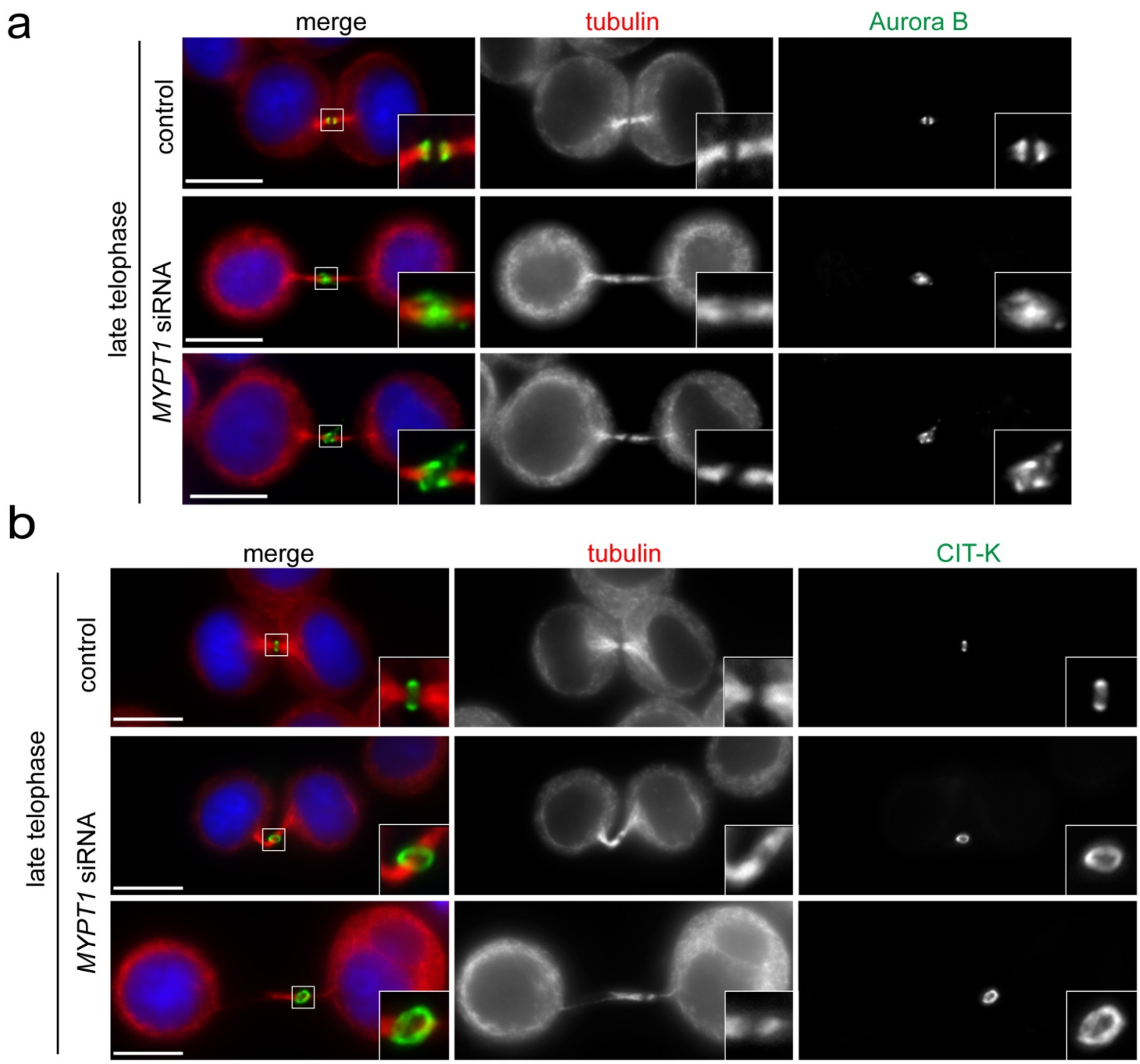


**Supplementary Fig. S2. Analysis of cytokinesis defects after MYPT1 depletion in HeLa S3 cells**. (a-b) HeLa S3 cells were treated with siRNAs directed against either a random sequence (control) or *MYPT1* and after 48 hours were fixed and stained to detect DNA (blue in the merged panels), tubulin, and either Aurora B (a) or CIT-K (b). DNA condensation and the shape and thickness of microtubule bundles at the intercellular bridge were used as criteria to stage telophase cells. Insets show a 3x magnification of the midbody. Bars, 10 µm.

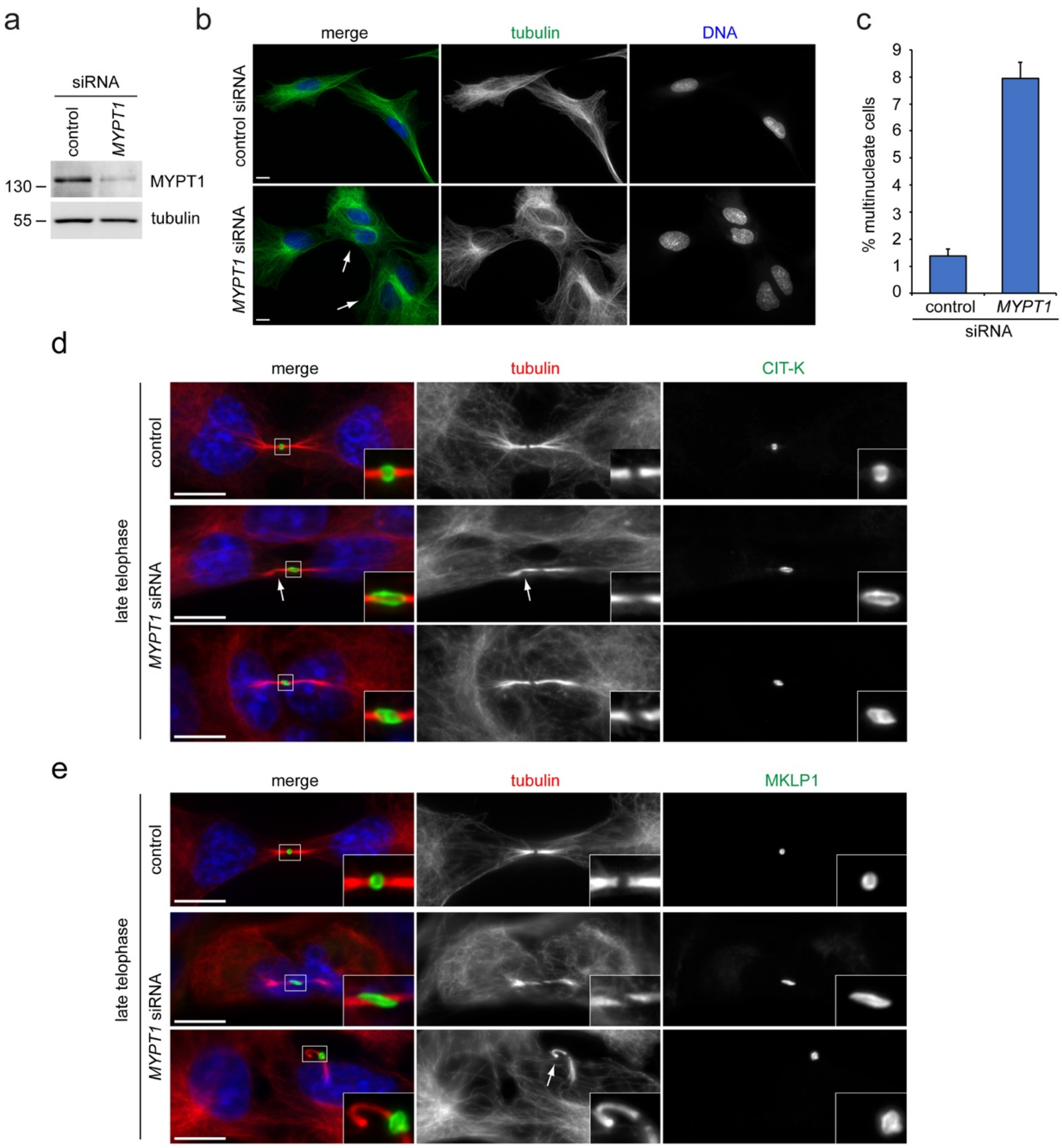


**Supplementary Fig. S3. Analysis of cytokinesis defects after MYPT1 depletion in RPE-1 cells**. (a) RPE-1 cells were treated with siRNAs directed against either a random sequence (control) or MYPT1 and after 48 h proteins were extracted and analyzed by western blot to detect the indicated proteins. The numbers on the left indicate the sizes in kDa of the molecular mass marker. (b) RPE-1 cells were treated with siRNAs directed against either a random sequence (control) or *MYPT1* and after 48 hours were fixed and stained to detect DNA and tubulin. The arrows indicate multinucleate cells. Bars, 10 µm. (c) Quantification of multinucleate cells obtained after control or MYPT1 siRNA. More than 700 cells were counted in each experiment, n=3. Bars indicate standard errors. (d-e) RPE-1 cells were treated with control or *MYPT1* siRNAs and after 48 hours were fixed and stained to detect the indicated epitopes and DNA (blue in the merged panels). DNA condensation and the shape and thickness of microtubule bundles at the intercellular bridge were used as criteria to stage telophase cells. Insets show a 3x magnification of the midbody. The arrows mark a bent (d) and a broken (e) central spindle, phenotypes that are very similar to those observed in HeLa cells (Fig. 6). Bars, 10 µm.

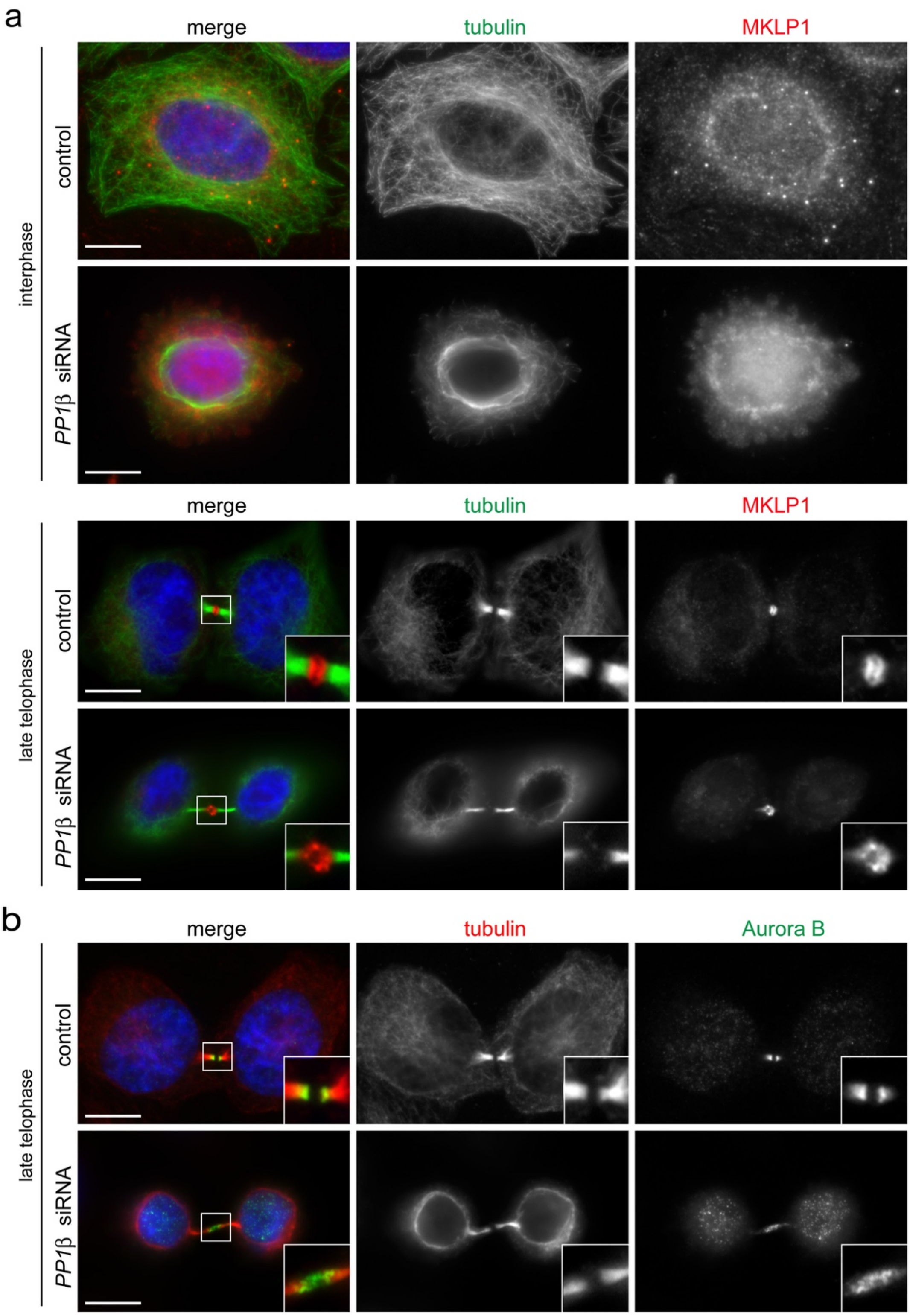


**Supplementary Fig. S4. *PP1β* siRNA phenocopies MYPT1 depletion**. (a-b) HeLa Kyoto cells were treated with siRNAs directed against either a random sequence (control) or *PP1β* and after 48 hours were fixed and stained to detect DNA (blue in the merged panels), tubulin and either MKLP1 (a) or Aurora B (b). DNA condensation and the shape and thickness of microtubule bundles at the intercellular bridge were used as criteria to stage telophase cells. Insets show a 3x magnification of the midbody. Bars, 10 µm.

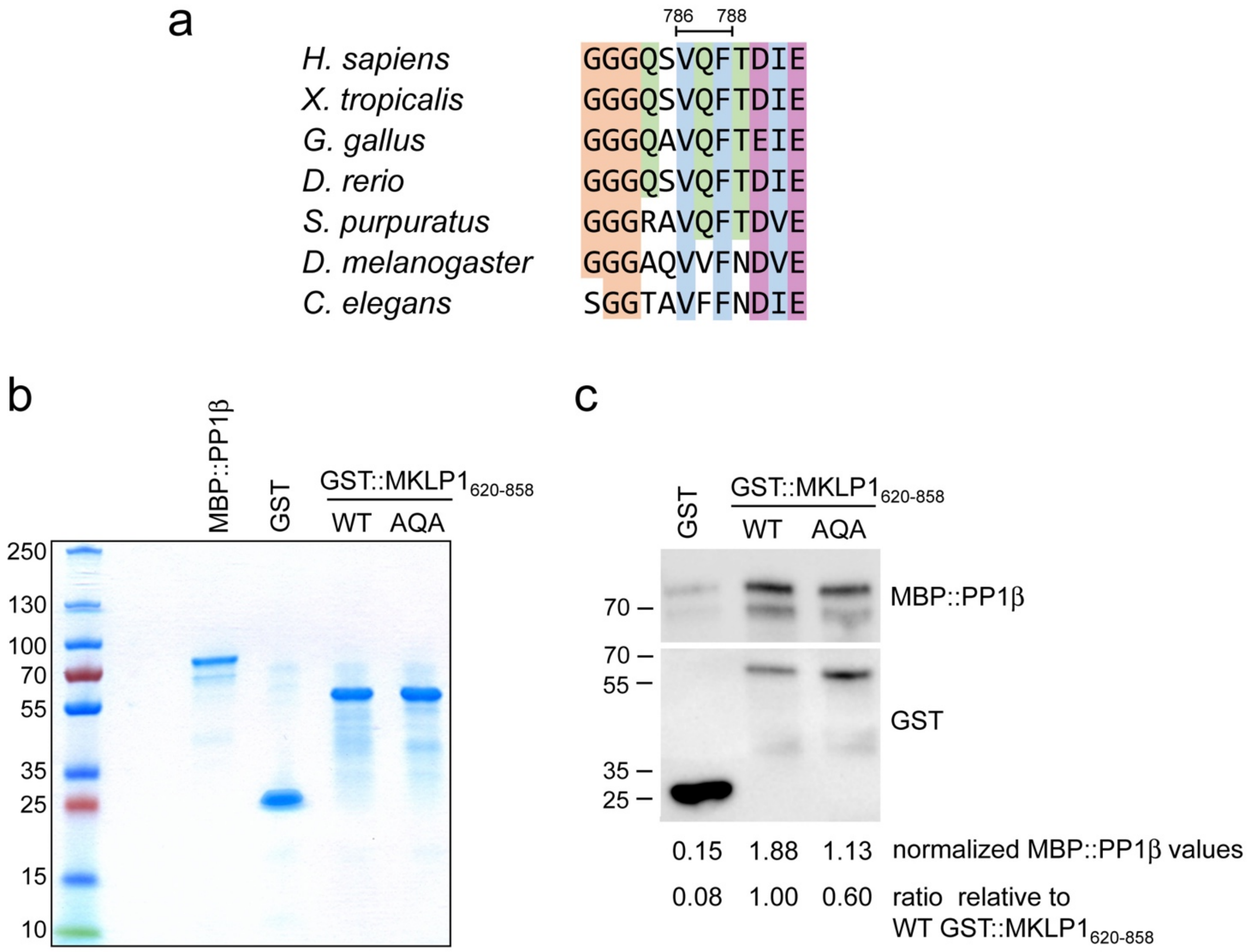


**Supplementary Fig. S5. PP1β interacts with a conserved binding site in the MKLP1 C-terminus.** (a) The amino acid sequences containing the PP1β binding site (aa 786-788) of human MKLP1 and its orthologues in other vertebrate and invertebrate species were aligned using the Muscle algorithm[1]. Amino acids are colored according to their chemical properties. (b) Coomassie-stained gel showing the purified proteins used in the GST pull down assay. (c) GST-tagged wild type and AQA mutant MKLP1 fragments (aa 620-858) were expressed and purified in bacteria and then employed in a pull down assay with MBP:: PP1β purified from yeast. Proteins were then analyzed by Western blot using antibodies against MBP (top) and GST (bottom). At the bottom are shown the values of MBP:: PP1β normalized against their respective GST baits and the ratio of these values relative to WT GST:: $MKLP1_{620-858}$.

**Supplementary Table S1. List of the proteins showing differential abundance at the midbody after CIT-K depletion.** Proteins that were significantly less or more abundant (*p* value < 0.01) in CIT-K siRNA midbodies are listed. The corresponding normalized logarithmic ratios and corrected p-values from experiment 1 and experiment 2 are shown. Proteins shaded in blue were underrepresented (negative logarithmic ratios) after CIT-K depletion. Proteins shaded in red were overrepresented (positive logarithmic ratios).

| Gene | Protein | $log_2$ ratio exp. 1 | $log_2$ ratio exp. 2 | *p*-value exp. 1 | *p*-value exp. 2 |
|---|---|---|---|---|---|
| CIT-K | Citron kinase | -2.69 | -2.47 | $2x10^{-45}$ | $1.4x10^{-25}$ |
| PC | Pyruvate carboxylase, mitochondrial | -0.80 | -0.86 | $3x10^{-9}$ | $3.9x10^{-4}$ |
| ASS1 | Argininosuccinate synthase | -0.52 | -0.63 | $7.6x10^{-5}$ | $5.5x10^{-4}$ |
| FLNB | Filamin B | -0.50 | -0.56 | $1.2x10^{-4}$ | $2.1x10^{-3}$ |
| TIMM44 | Mitochondrial import inner membrane translocase subunit TIM44 | -0.40 | -1.04 | $1.4x10^{-3}$ | $1.82x10^{-5}$ |
| PPIB | Peptidyl-prolyl cis-trans isomerase B | -0.37 | -0.57 | $3.1x10^{-3}$ | $1.8x10^{-3}$ |
| AURKA | Aurora kinase A | 0.45 | 0.69 | $5x10^{-3}$ | $2.6x10^{-4}$ |
| CKAP2 | Cytoskeleton-associated protein 2 | 0.45 | 0.67 | $5x10^{-3}$ | $3.7x10^{-4}$ |
| PTPRF | Receptor-type tyrosine-protein phosphatase F | 0.62 | 1.19 | $2.3x10^{-3}$ | $9.71x10^{-10}$ |
| TPX2 | Targeting protein for Xklp2 | 0.71 | 0.75 | $4.3x10^{-6}$ | $6.65x10^{-5}$ |
| HNRNPH3 | Heterogeneous nuclear ribonucleoprotein H3 | 0.83 | 0.97 | $3.2x10^{-5}$ | $5.47x10^{-7}$ |
| KIFC1 | Kinesin-like protein KIFC1 | 0.87 | 0.55 | $1.5x10^{-5}$ | $3.3x10^{-3}$ |

**Supplementary Table S2. List of the serine/threonine phosphatases present in the midbody interactome**. The baits are listed in the top row and the phosphatases in the far left column. Protein names are according to the UniProt database. Member of the PP1 family are shaded in green, while members of the PP2 family are shaded in red. The Mascot score (Score) and number of peptides (Pept) identified in each AP-MS experiment are indicated. The far-right column indicates whether the phosphatase was identified in the midbody proteome.

| Baits | Anillin | Aurora B | CHMP4B | CHMP4C | CIT-K | ECT2 | KIF14 | KIF20A | KIF23/ MKLP1 | PRC1 | Proteome |
|---|---|---|---|---|---|---|---|---|---|---|---|
| **PPP1CA** | Score 183<br>Pept 5 | | | Score 244<br>Pept 6 | Score 1392<br>Pept 21 | Score 400<br>Pept 9 | Score 658<br>Pept 10 | Score 303<br>Pept 8 | Score 75<br>Pept 3 | Score 1004<br>Pept 16 | Yes |
| **PPP1CB** | | | | | Score 1312<br>Pept 19 | Score 342<br>Pept 8 | Score 85<br>Pept 1 | Score 253<br>Pept 6 | | Score 963<br>Pept 14 | No |
| **PPP1CC** | Score 267<br>Pept 9 | Score 117<br>Pept 1 | Score 302<br>Pept 6 | | Score 1355<br>Pept 20 | Score 338<br>Pept 8 | Score 690<br>Pept 12 | Score 407<br>Pept 6 | Score 68<br>Pept 2 | Score 1067<br>Pept 19 | Yes |
| **PPP1R9A** | Score 235<br>Pept 3 | | Score 120<br>Pept 2 | Score 83<br>Pept 3 | | Score 254<br>Pept 4 | Score 67<br>Pept 1 | Score 48<br>Pept 1 | | Score 78<br>Pept 1 | No |
| **PPP1R9B** | Score 237<br>Pept 5 | | Score 285<br>Pept 6 | | | Score 340<br>Pept 7 | Score 273<br>Pept 5 | Score 94<br>Pept 1 | Score 98<br>Pept 2 | | No |
| **PPP1R12A (MYPT1)** | Score 185<br>Pept 2 | | Score 197<br>Pept 3 | Score 60<br>Pept 5 | Score 572<br>Pept 6 | Score 363<br>Pept 11 | Score 855<br>Pept 15 | Score 152<br>Pept 5 | | Score 564<br>Pept 13 | Yes |
| **PPP1R12B** | Score 87<br>Pept 1 | | | | | | Score 83<br>Pept 1 | | | | No |
| **PPP1R12C** | | | | | | | Score 346<br>Pept 4 | Score 87<br>Pept 1 | | Score 56<br>Pept 1 | No |
| **PPP1R18** | Score 96<br>Pept 1 | | Score 128<br>Pept 1 | | Score 85<br>Pept 1 | Score 232<br>Pept 4 | Score 400<br>Pept 8 | Score 201<br>Pept 7 | | | No |
| **PPP1R13L** | | | | | | | Score 101<br>Pept 2 | Score 222<br>Pept 2 | | | Yes |
| **PPP2CA** | | | | Score 38<br>Pept 1 | | | | | | | No |
| **PPP2R5A** | | | | Score 24<br>Pept 3 | Score 107<br>Pept 2 | | Score 34<br>Pept 1 | | | Score 98<br>Pept 2 | Yes |
| **PPP2R1B** | | | | | | | | | | Score 46<br>Pept 1 | Yes |
| **PPP3CA** | | | Score 61<br>Pept 1 | | | Score 63<br>Pept 1 | | | | | No |
| **MPRIP** | Score 1358<br>Pept 23 | | Score 390<br>Pept 6 | Score 43<br>Pept 3 | Score 66<br>Pept 1 | Score 179<br>Pept 6 | Score 528<br>Pept 10 | Score 432<br>Pept 5 | Score 85<br>Pept 2 | | No |
| **CDC25A** | | Score 48<br>Pept 9 | | | Score 39<br>Pept 3 | | | | | Score 35<br>Pept 1 | No |
| **PGAM5** | Score 359<br>Pept 11 | | Score 75<br>Pept 2 | Score 108<br>Pept 4 | Score 61<br>Pept 4 | Score 74<br>Pept 3 | Score 430<br>Pept 13 | Score 372<br>Pept 13 | Score 171<br>Pept 6 | Score 101<br>Pept 2 | Yes |

### Supplementary Video Captions

**Supplementary Video S1.**

This movie shows the different focal planes (z sections at 0.25 μm step size) of the tubulin immuno-staining of the late telophase control cell depicted in Fig. S5A (left bottom panels).

**Supplementary Video S2.**

This movie shows the different focal planes (z sections at 0.25 μm step size) of the tubulin immuno-staining of the late telophase *MYPT1* siRNA cell depicted in Fig. S5A (right bottom panels).

**Supplementary Video S3.**

This movie shows the 3D reconstruction of the tubulin immuno-staining of the late telophase control cell depicted in Fig. S5A (left bottom panels).

**Supplementary Video S4.**

This movie shows the 3D reconstruction of the tubulin immuno-staining of the late telophase *MYPT1* siRNA cell depicted in Fig. S5A (right bottom panels).

**Supplementary Video S5.**

This movie shows GFP::tubulin and histone H2B::mCherry dynamics in a control cell that successfully completed cytokinesis; see Fig. 3A for details. Cells were treated with dsRNA directed against a random sequence (control) for 30 hours before start filming. See Figure 3A for details. All sequences were captured at 2 min intervals. Playback rate is 5 frames per second (FPS).

**Supplementary Video S6.**

This movie shows GFP::tubulin and histone H2B::mCherry dynamics in an *MYPT1* siRNA cell. Note the long and thin central spindle that snaps after more than 2 hours after anaphase onset; see Fig. 3A for details. Cells were treated with dsRNA directed against *MYPT1* for 30 hours before start filming. All sequences were captured at 2 min intervals. Playback rate is 5 frames per second (FPS).

**Supplementary Video S7.**

This movie shows GFP::tubulin and histone H2B::mCherry dynamics in an *MYPT1* siRNA cell. In this cell, a long and thin central spindle forms after completion of furrow ingression, but there is no abscission and cytokinesis ultimately fails forming a single binucleate cell. Cells were treated with dsRNA directed against *MYPT1* for 30 hours before start filming. All sequences were captured at 2 min intervals. Playback rate is 5 frames per second (FPS).

**Supplementary Video S8.**

This movie shows GFP::tubulin and histone H2B::mCherry dynamics in an *MYPT1* siRNA cell. This cell shows abnormal contractility during furrow ingression, fails to properly assemble the central spindle and consequently cytokinesis fails prior to midbody formation. Cells were treated with dsRNA directed against *MYPT1* for 30 hours before start filming. All sequences were captured at 2 min intervals. Playback rate is 5 frames per second (FPS).

### Supplementary Data Captions

**Supplementary Data S1. (separate file)**

Excel file listing the proteins identified by MS from the pull downs of CIT-K::AcGFP expressing cells at different cell cycle stages (see Fig. 1A, B). Each worksheet contains the results for a specific cell cycle stage, i.e., S phase, metaphase and telophase. Non-specific binding proteins were removed by filtering each dataset against the corresponding dataset obtained from AP-MS experiments using HeLa cells expressing GFP alone at the same cell cycle stage (see Materials and Methods).

**Supplementary Data S2. (separate file)**

Excel file listing the proteins identified in the SILAC experiments (Fig. 1C to E) and the relative quantification analysis described in Materials and Methods.

**Supplementary Data S3. (separate file)**

Excel file listing the proteins identified by MS from the pull downs of telophase HeLa cells expressing the ten different baits indicated in Table S2. Each worksheet contains the results for a specific bait. Non-specific binding proteins were removed by filtering each dataset against the corresponding dataset obtained from AP-MS experiments using telophase HeLa cells expressing GFP alone (see Materials and Methods). The MS data for Flag::CHMP4C are from our previous study[2].

**Supplementary Data S4. (separate file)**

Excel file listing the proteins identified in the midbody proteome and interactome along with their respective gene names and GO terms. The proteins shared between the two datasets and the ones specific for each dataset are shown in separate worksheets.

**Supplementary Data S5. (separate file)**

Excel file showing the results of the GO enrichment analyses of the midbody proteome and of the midbody interactome in two separate worksheets.

### References


1 Edgar, R. C. MUSCLE: multiple sequence alignment with high accuracy and high throughput. *Nucleic Acids Res* **32**, 1792-1797, doi:10.1093/nar/gkh340 (2004).

2 Capalbo, L. *et al.* Coordinated regulation of the ESCRT-III component CHMP4C by the chromosomal passenger complex and centralspindlin during cytokinesis. *Open Biol* **6**, doi:10.1098/rsob.160248 (2016).
